# Particle Tracking facilitates real time Motion Compensation in 2D or 3D two-photon imaging of neuronal activity

**DOI:** 10.1101/112284

**Authors:** Samira Aghayee, Zach Bowen, Daniel E. Winkowski, Matt Harrington, Patrick O. Kanold, Wolfgang Losert

## Abstract

The application of 2-photon laser scanning microscopy (TPLSM) techniques to measure the dynamics of cellular calcium signals in populations of neurons is an extremely powerful technique for characterizing neural activity within the central nervous system. The use of TPLSM on awake and behaving subjects promises new insights into how neural circuit elements cooperatively interact to form sensory perceptions and generate behavior. A major challenge in imaging such preparations is animal and tissue movement, which leads to shifts in the imagine location (jitter). Although there are surgical and technical approaches to minimize brain motion under these conditions, it is generally unavoidable. The presence of image motion can lead to artifacts, especially since quantification of TPLSM images involves analysis of fluctuations in fluorescence intensities for each neuron, determined from small regions of interest (ROIs) Here, we validated a new motion correction approach to compensate for motion of TPLSM images in the superficial layers of auditory cortex of awake mice. We use a nominally uniform fluorescent signal as a secondary signal to complement the dynamic signals from genetically encoded calcium indicators. We tested motion correction for single plane time lapse imaging a; well as multiplane (i.e. volume) time lapse imaging of cortical tissue. Our procedure of motion compensation relies on locating the brightest neurons and tracking their positions over time using established techniques of particle finding and tracking. The performance of our techniques both for 2D (single plane) and 3D (volume) image sequences is comparable to established techniques in its ability to suppress false neural signals. Object tracking based motion compensation thus offers an alternative approach for motion compensation, one that is well suited for real time feedback control and for analysis of tissue distortions.

## Background and introduction

Two-photon laser scanning microscopy (TPLSM) of neuronal activity using Ca^2+^ indicators is a powerful new approach that is allowing us to analyze the inner workings of the brain at the level of single cells for large groups of neurons within an awake behaving organism(Packer et al., 2012). While electrophysiology approaches allow for very accurate measurements of the neuronal circuit a few neurons at a time, TPLSM allows for the simultaneous observation of hundreds and thousands of neurons (Peron et al., 2015) and thus yields information on the collective behavior of groups of neurons. Understanding the collective behavior of neurons is essential for understanding how the brain processes information and encodes memory (Beggs and Plenz, 2003; Kinouchi and Copelli, 2006; Rämö et al., 2007; Shew et al., 2009; Shew et al., 2011; Yang et al., 2012; Shew and Plenz, 2013). The essence of these analyses is that behaviorally relevant information in the brain does not reside in the firing events of individual neurons, but instead in the collective firing behavior of groups of neurons.

Measurements of collective dynamics require accurate detection of neuronal activity of tens to thousands of individual neurons. Since detection errors grow exponentially with the size of the observed neuronal population, detection inaccuracies would lead to spurious measurements and conclusions. For example, with 97% accuracy for identification of a single neuronal spike one can only achieve 74% accuracy for identifying simultaneous activity of 10 neurons. Thus, to detect synchronous activity of 100 neurons with more than 95% confidence the detection of single neuron events has to be made with a close to 100% accuracy.

A precise spike inference is particularly challenging in two-photon Ca^2+^ imaging data where neuronal activity is inferred by brightness fluctuations of imaged pixels within the regions of interest (ROI). Current indicators such as GCaMP6 (Chen et al., 2013) are expressed in the cytoplasm of the neuron, forming a ring surrounding the nucleus, thus when imaging a large population of neurons, relatively few pixels contribute to the cellular signal. Displacement of the neuronal somata over time from the defined ROIs, can therefore perturb the inferred activity. Hence, it is essential to accurately correct for motion of the image plane that can be particularly large in awake behaving preparation before extracting neuronal activity. Moreover, to facilitate closed loop experiments, motion correction has to operate fast enough to extract neuronal activity in real time.

To address these needs, we have developed a suite of analytical tools to quantify brain motion that adapts sub-pixel accuracy object-tracking tools from the field of soft-matter physics. We use these tools to characterize brain motion and to compensate for it with sub-pixel resolution at close to real time speed. In addition, we compare the performance of our tools to established algorithms that are used widely for motion correction. Specifically, we introduce a particle-tracking based image analysis pipeline capable of significantly reducing in-plane motion, which yields more accurate spike inference measurements. The algorithm operates close to real time, and is amenable to extensions for 3D motion compensation.

### Methods

#### Animals

All procedures were approved by the University of Maryland Institutional Animal Care and Use Committee.

Adult wild type (C57) mice (>P40, range P40 to P100 at the time of experiments) of both genders underwent a single aseptic surgical procedure in which they received an implant of a titanium headplate, intracortical injections of genetically encoded calcium indicators, and chronic cranial windows (Goldey et al., 2014). The titanium headplate design was a modified version of headplate presented by Guo et al. (Guo et al., 2014) to allow access to auditory cortex. Adeno-associated virus (AAV-syn-mRuby2-GCaMP6s(Rose et al., 2016) was obtained from UPenn Vector Core and injected into several locations (~30nL per site; ~300-350μm from pial surface; 3-6 sites) in the auditory cortex using a Nanoject system (Drummond). The craniotomy was sealed with a chronic cranial window (Goldey et al., 2014). The entire implant except for the window over the auditory cortex was covered in black dental acrylic. At the conclusion of the procedure, the animals were given a subcutaneous injection of meloxicam as an analgesic. After complete recovery from surgery (>1-2 weeks), animals were acclimatized to the restraint system over several sessions. After 1 week of acclimatization, 2-photon imaging experiments commenced.

#### Imaging

For 2-photon imaging, we used a scanning microscope (Bergamo II series, B248, Thorlabs) coupled to a pulsed femtosecond Ti:Sapphire 2-photon laser with dispersion compensation (Vision S, Coherent). The microscope was controlled by ThorImageLS software. The laser was tuned to λ = 940 nm in order to simultaneously excite GCaMP6s and mRuby2. Red and green signals were collected through a 16× 0.8 NA microscope objective (Nikon). Emitted photons were directed through 525/50-25 (green) and 607/70-25 (red) band pass filters onto cooled GaAsP photomultiplier tubes (Hammamtsu). The field of view was 370 × 370 μm. Imaging frames of 512×512 pixels (pixel size 0.72 μm) were acquired at 30Hz by bidirectional scanning of an 8 KHz resonant scanner. Beam turnarounds at the edges of the image were blanked with a Pockels cell. The average power for imaging was <70 mW, measured at the sample. For single plane imaging, imaging sites were ~150-270 μm from the pia surface.

For volume scanning, axial motion was controlled with a piezo collar (Physik Instrument) attached to the microscope body. The microscope objective was moved smoothly through through a z-distance of ~50 μm across 11 imaging planes, two frames as the objective returned to the starting position (i.e., flyback frames) were discarded. All images were acquired at 352 × 352 pixels (253 μm × 253 μm). The inclusion of multiple z-planes reduced our temporal resolution to ~3 frames per second.

#### Image Analysis Algorithm

##### Cell finding algorithm

We use a cell tracking approach for motion compensation. To extract the position of cells in the invariant channel over background we use bandpass filtering and peak finding. Figure 1A shows a sample image of low signal to noise ratio. In this example we use a 16x objective and image 512 × 512 pixels (at 0.7215 μm per pixel) such that a cell diameter is about 15 pixels. To enhance the signal to noise ratio, the image is bandpassed with a lower threshold of 4 pixels to remove pixel scale noise and an upper threshold comparable to (typically slightly smaller than) the cell size (13 pixels). The resulting image is shown as Figure1B. On this filtered image we use automatic peak finding routines(Crocker and Grier, 1996) to locate the position and size of each peak in the image above a threshold. The center is interpolated based on the brightness of all pixels in the bandpassed image within a user-defined region of radius R independent of the bandpass thresholds, but it should not be chosen similar to the upper bandpass value.

**Figure 1.**
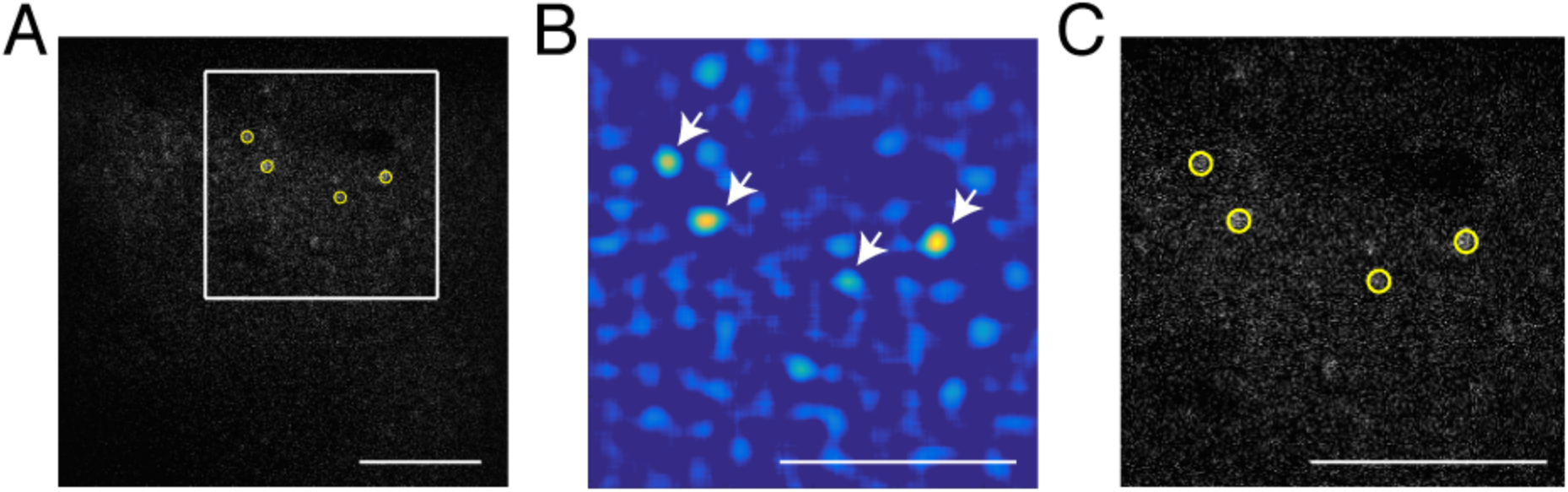
Particle finding via bandpass filtering. The scale bar is 100 μm. a) Typical frame acquired by TPSM. The low signal to noise ratio is in most cases unavoidable due to high speed scanning. b) Bandpassed image of the same frame with lower and higher bandpass of respectively 4 and 13 pixels. c) The four brightest cells extracted by the algorithm (yellow circles) Scale bar is 100 microns.

##### Cell Tracking algorithm

Once cell positions are determined in each frame, individual cells are mapped from frame to frame to enable unique identification of each cell. This is accomplished by mapping cells from one frame to the next in such a way that the cumulative square displacement between frames is a minimum value,

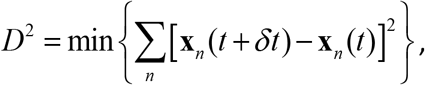

where *X_n_* corresponds to the positions of the n^th^ cell in a particular frame. The tracking software can also be adjusted with a memory parameter to account for the possibility that some cells may not be visible enough to be located in a subset of the frames. Cells are then “remembered” for the subset of frames in which they are not visible. This mapping yields trajectories for each cell that could be mapped through the image sequence.

##### Motion Correction

From the shift in all measured trajectories in each pair of frames we then infer the overall shift and rotation of each frame. While the shift in the positions of all tracked neurons does not perfectly fit a rotation and translation (local distortions are also possible) we compute the rigid body transformation (translation and rotation) that minimizes residual displacements from one frame to the next using the Kabsch algorithm (Kabsch, 1976; 1978). These rigid body transformations are calculated for all frames to carry out motion compensation for the whole image sequence.

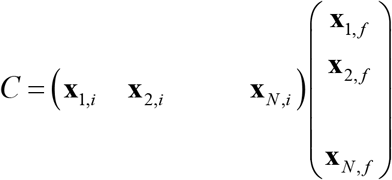

Here i represents the positions in the frame (source) and f the average positions we want to correct towards (target)

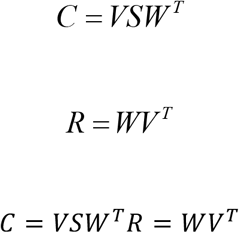

##### Validation of Subpixel motion correction

Subpixel precision can be achieved via peak finding, but only if each peak spreads over a sufficient number of bright pixels. If subpixel precision is achieved, the jitter amount and cell motion, when measured in pixels, should equally likely fall anywhere between integer pixel values. Thus the residuals should have a random distribution between 0 and 1. Conversely, if peak finding and motion correction only achieved pixel resolution, the distributions would peak around both 0 and 1. Examples of uniform histograms for the residual values of both cell motion and frame jitter are shown in Figure 2.

**Figure 2.**
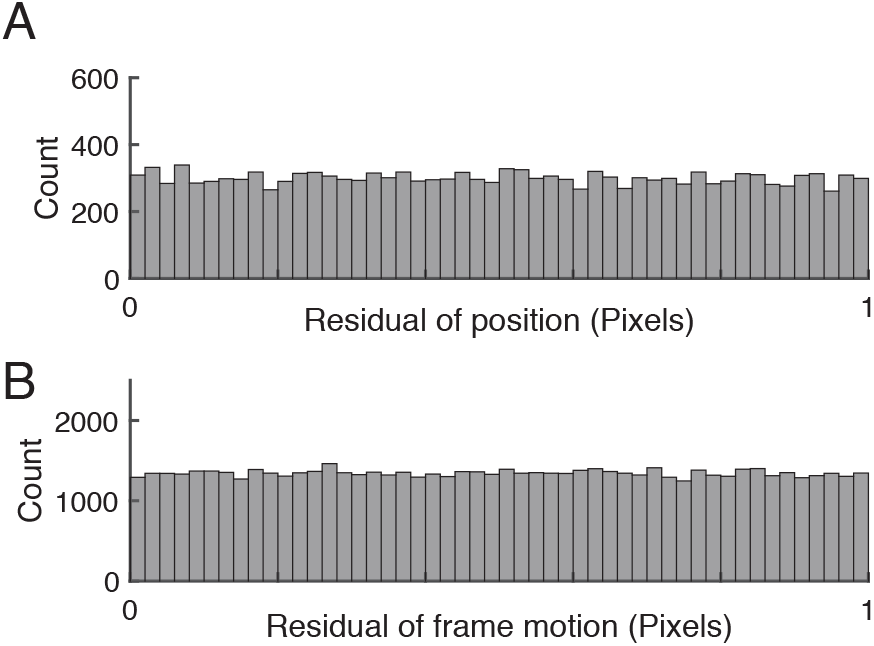
Validation of subpixel resolution particle finding and motion compensation. a) Distribution of the residual cell position after extraction for 67148 cells extracted from 15000 frames b) Distribution of the frame-to-frame residual displacement. The uniform distributions indicate sub-pixel resolution of both cell tracking and motion compensation.

### Results

We quantify brain motion using object-tracking methods adapted from the field of soft-matter physics outlined above. Fast, robust particle tracking algorithms have been first introduced two decades ago (Crocker and Grier, 1996) and refined extensively since then by others (Blair and Dufresne, 2008) and us (Losert et al., 2000). The particle tracking approach, described in the Methods section in detail, takes advantage of the fact that neuronal cell bodies are roughly spheroidal and resemble each other in size and shape. Our approach works best with a signal that is independent of neuronal activity and thus under ideal circumstances would appear at uniform brightness. This is achieved by transfecting neurons with constructs that besides supplying a Ca^2+^ indicator also label the nucleus or cytosol with activity independent label, e.g. mRuby2 or others (Peron et al., 2015; Rose et al., 2016). We benchmark our algorithm using image sequences obtained from the primary auditory cortex in awake mice using GCamp6s for Ca^2+^ and mRuby2 as a somatic marker (AAV-syn-mRuby2-GCaMP6s).

Applying peak finding and particle tracking, detailed in the methods section, yields the motion of each neuron independently. The time dependent position of a single neuron is shown in Figure 3A. Since tracks of nearby neurons look very similar (Figure 3B) it is reasonable to use the tracks for motion compensation as described above. We compute the translational and rotational shift of an image from tracks of more than two points (N>2). Two reference points, though sufficient in principle, are not enough given the uncertainty and errors in both imaging and processing. Since peak finding increases in accuracy with increased neuronal brightness, we use the brightest N tracked neurons for the motion compensation algorithm outlined in the Methods section. The power spectrum of jitter, shown in Figure 3C, exhibits notable peaks on top of a regular noise spectrum at frequencies of 7.7Hz and 9.7Hz comparable to the expected heartbeat of a mouse. Correcting the position of each point for this jitter yields close to stationary points, with only very weak residual fluctuations, as shown in Fig 3D.

**Figure 3.**
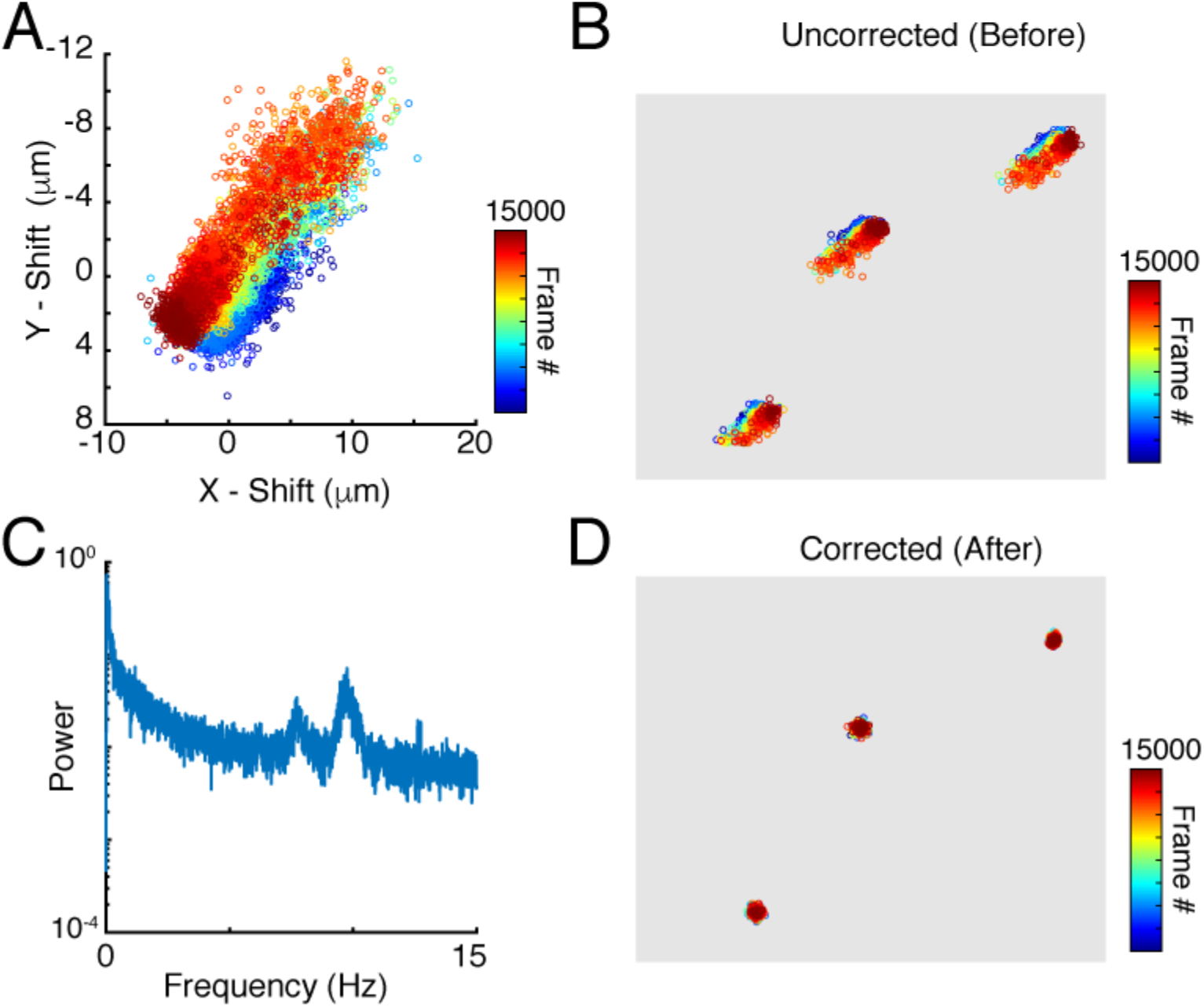
Particle tracking and residual cell motion a) Tracks of a single neuron in time. b) Tracks for three neurons highlight the similarities in trajectories c) Power spectrum of image jitter, d) Tracks after motion compensation.

When benchmarking for processing speed, our algorithm yields analysis of 15000 images in 779.8 seconds or 19.2 frames per second on a six core 3.5 GHz Intel Xeon Mac with OSX and 64Gb of RAM.

The next step is to use the motion compensated images to identify neuronal Regions of Interest (ROI) to be used for measurement of time traces of activity. Motion compensation yields a sharp averaged image (Figure 4B) compared to the uncorrected average (Figure 4A). While a single frame is so noisy that peak finding only captures the brightest neurons accurately (Figure 1), peak finding on the averaged image reliably yields most of the cells visible by eye (Figure 4C). These brightness peaks provide automated input for established algorithms to identify regions of interest (ROI) around each cell center, and to trace the image intensity in the ROI over time.

**Figure 4.**
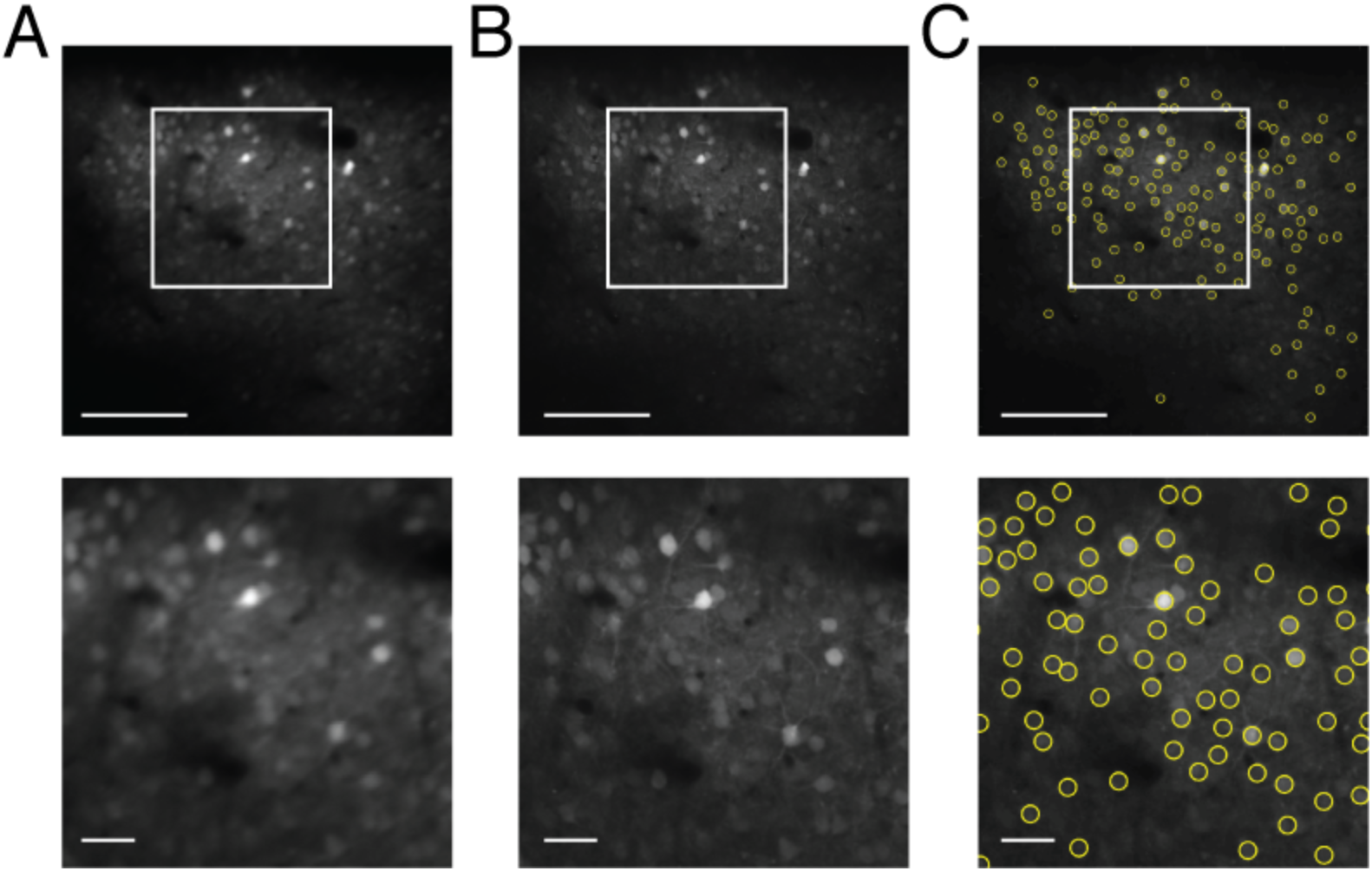
Cell finding from motion compensated averaged images. The scale bars are 100μm (top) and 20μm (bottom) A) Averaged image for the unregistered image sequence B) Averaged image after motion compensation C) Automated cell finding.

To measure motion compensation quality, we take advantage of the nominally uniform brightness of the second (in this case red) fluorophore with time. Therefore the fluorescence intensity in each of the motion compensated cell ROI should be constant. Fluctuations are indicative of both inaccuracies in motion compensation and measurement noise, since individual images are noisy as seen in Figure 1. We first measure the time sequence of these fluctuations using the z-score as a normalized measure. Figure 5 shows the z-score of one representative cell as a function of time for the original image sequence, after full image registration with TurboReg (Thevenaz et al., 1998), and after tracking motion compensation. The unregistered image sequence has several sudden jumps in z-score indicative of sudden large changes in position, as well as slow drift in z-score indicative of slower shifts in the imaging location. Motion events (blue arrows) in particular may be misidentified as neuronal activity in spike inference algorithms. Both Turboreg and particle tracking based motion compensation eliminate these artifacts. For all tracked neurons, the z-scores obtained with both motion compensation methods are comparable for both the second control fluorophore (Figure 5B) and for the calcium signal (Figure 5C).

**Figure 5.**
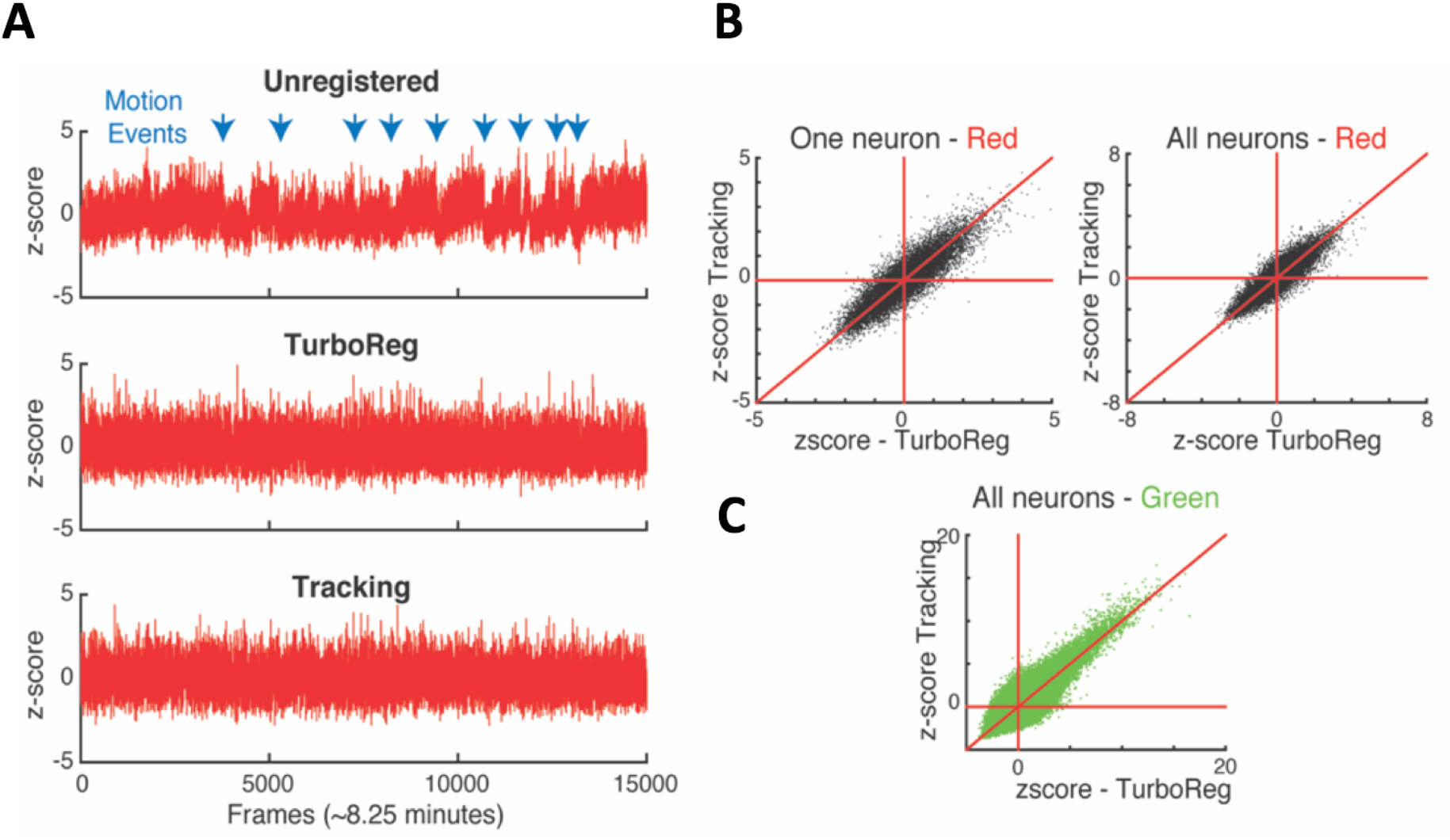
A: Comparison of normalized intensities (z-scores) of the reference channel vs time for one representative neuron. B: The z-scores obtained with tracking and turboreg based motion compensation match for one representative neuron, and all neurons. C: Comparison of z-scores in the calcium imaging channel (green) for tracking and turboreg corrected images.

To assess how the quality of motion compensation depends on tracking parameters and the number of tracked cells, we analyzed the same image sequence with multiple parameters, and with DFT for comparison. For a robust baseline comparison to established practice, we used manual cell identification from the averaged images for all cases. Figure 6A shows the intensity fluctuation in the reference signal (the static channel) measured from the four brightest cells (red) and for all 254 cells detectable in the averaged image (black). The tracking based analysis is robust to changes in parameters (for the number of neurons N between 4 and 10 and bandpass levels of 13 to 15 pixels, comparable to the size of a neuron), and comparable to the performance of DFT. Since the fluorescence from the control channel should be constant in time for each neuron, and at the same time the fluorophore uptake and thus fluorescence intensity levels vary significantly from neuron to neuron, we can determine whether fluctuations in measured intensity depend on the brightness of the neuron. Figure 6B shows that after motion compensation, the fluctuations decrease with increasing brightness of the cell. For unregistered images, on the other hand, fluctuations are independent of brightness. The dependence on brightness approximately follows an inverse square root dependence on intensity, with a small intensity offset, as expected for a noisy system.

**Figure 6.**
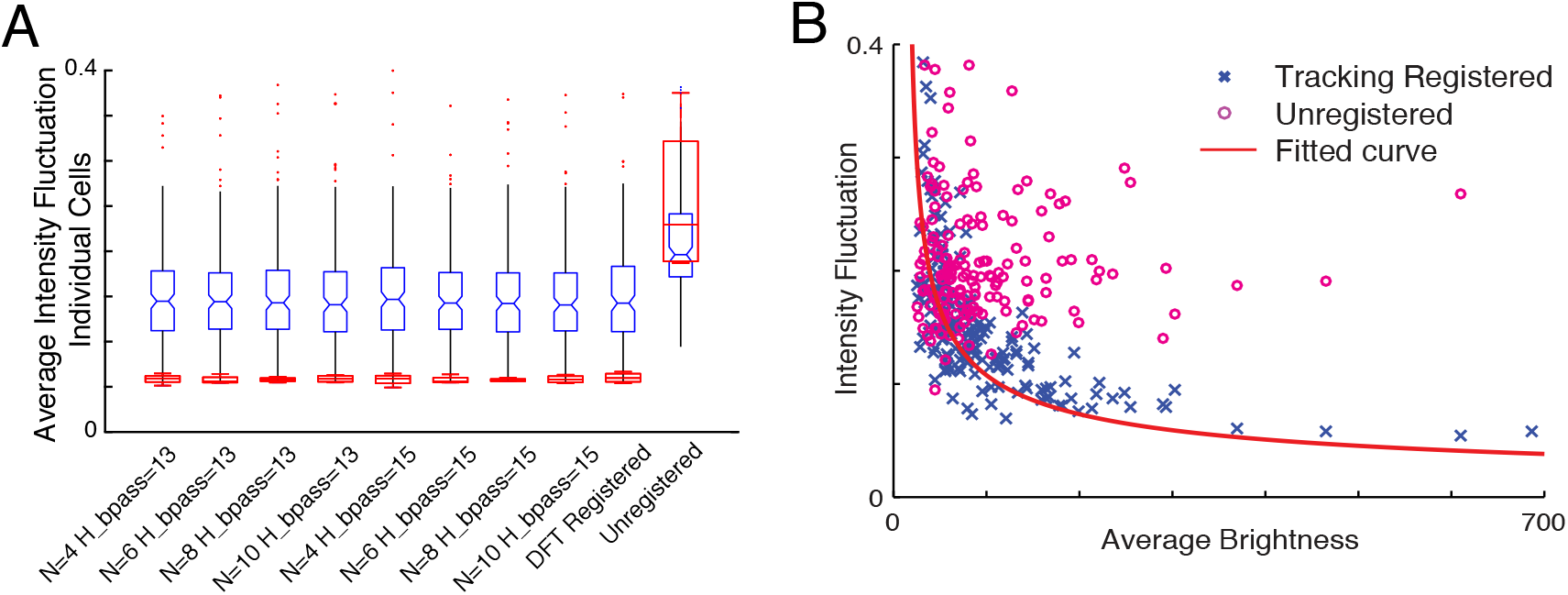
a) Comparison of intensity fluctuations in the static channel for different tracking parameters with the DFT corrected sequence and unregistered stack. Blue: all cells, Red: four brightest cells b) Intensity fluctuation versus average brightness of each cell in the static channel. The blue cross represents ROI extracted from the tracking registered stack while the circles in magenta are shows the ROI extracted from the unregistered stack. The red line shows the fitted curve to the registered ROIs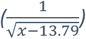

Finally, we compare whether our new tracking based motion compensation yields a good SNR of the Ca^2+^ signal that represents neuronal activity fluctuations in the GCamp6 channel. This measurement is particularly sensitive to the robustness of the algorithm since the SNR depends on peaks in brightness above a threshold as shown in Figure 7A. We find very similar, if not slightly higher, SNR for each neuron with our tracking based motion compensation approach when compared to Turboreg.

**Figure 7.**
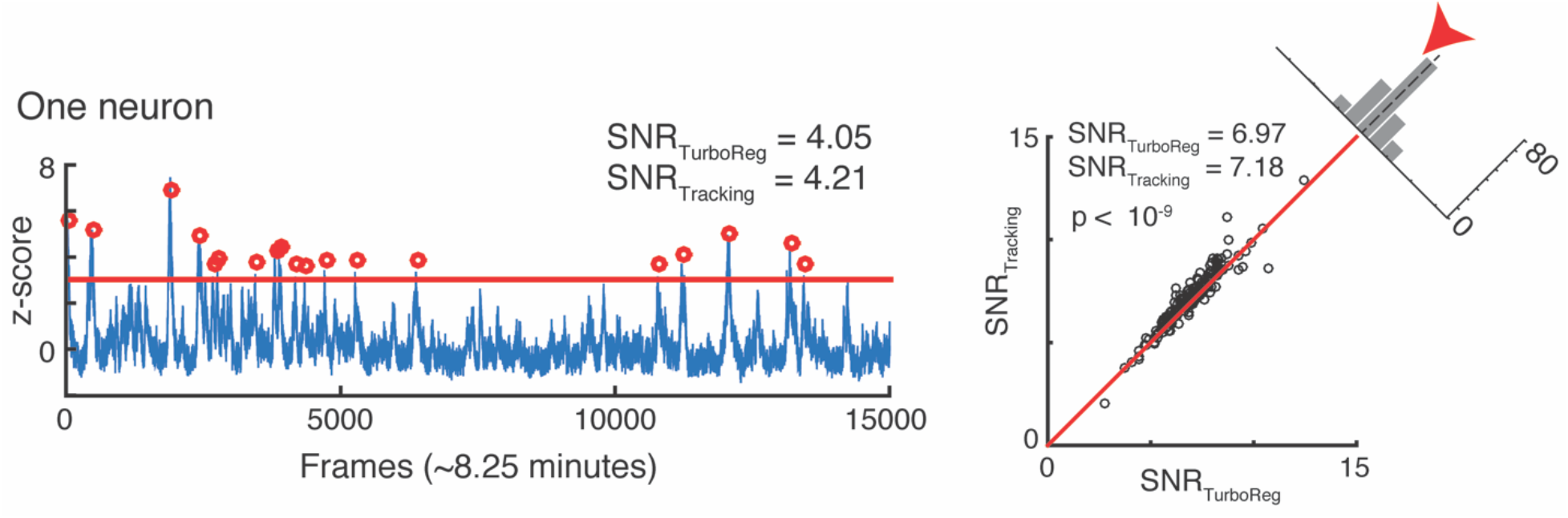
a) SNR calculation for one neuron b) SNR correlation of tracking registration vs TurboReg

As an added benefit, the tracking based approach to motion compensation also registers volume images. Volume imaging has become feasible due to the availability of faster scanning methods. Since Ca^2+^ signal has a time constant of 500ms (Chen et al., 2013) it is now possible to record neuronal activity in several z-regions with a frame rate of 30Hz without significantly diminishing acquisition concurrency. Volume imaging can allow for imaging of neurons in different layers, or to provide additional information about the same neurons. Since our in-plane motion compensation approach adjusts image positions to their time averaged position, it automatically yields images that are registered in plane across multiple planes. This is demonstrated in Figure 8, which shows, for 6 neurons in an image (Figure 8A) an overlay of the measured positions of the same cells visible in multiple imaging planes before motion compensation (Figure 8B) and after motion compensation carried out for each image plane independently (Figure 8C). Motion compensation to the average position yields a simple, yet powerful approach to create aligned 3D volume time lapse images. From such registered image stacks it is then possible to identify the average z-position of a neuron as well based on the z-dependent fluorescence intensity of the ROI around a peak, as shown in Figure 8D,E. Thus our approach can be extended to ultimately compensate for motion perpendicular to the imaging plane as well.

**Figure 8.**
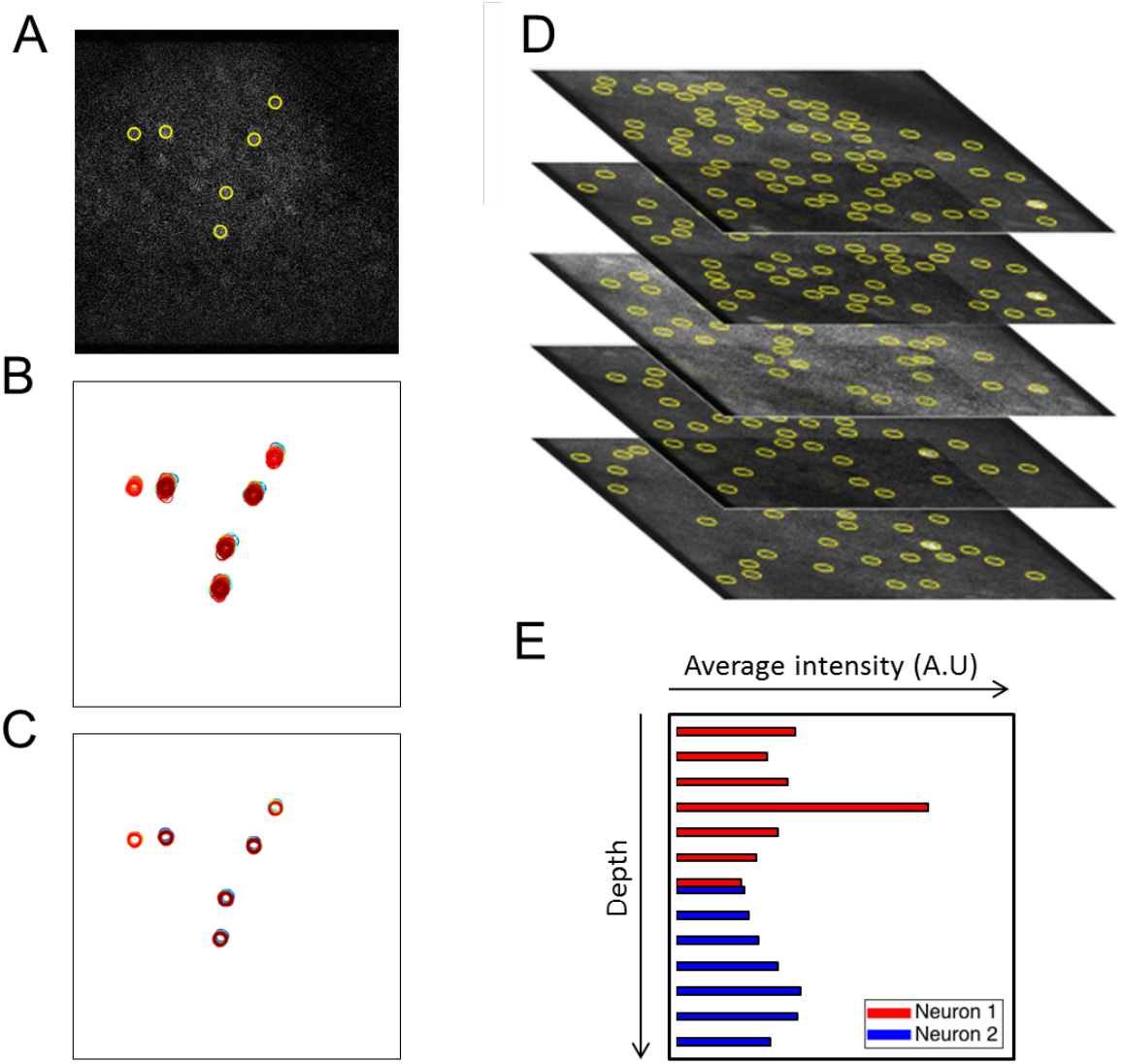
Extension of peak finding to three dimensions. a) Sample image with 6 extracted cells. Overlay of the six cells identified in all layers at all times (unregistered) c) Overlay of cell position after motion compensation d) Brightest cells are extracted independently from each z-plane of the volume data image. Extracted cells are tracked through data acquisition timepoints for every plane. e) Intensity of two sample cells in different Z planes.

## Discussion

We introduce peak finding and particle tracking as a new motion compensation approach that is fast and compatible with the low SNR nature present in TPLSM neural activity data. It works best, with a time invariant fluorescent signal, generated with a secondary fluorophore. One alternative may be to add rather than subtract intensities of the two channels of a FRET probe such as YC2.6 (Whitaker, 2010). Without optimization we achieve close to real time analysis speeds.

Our approach complements whole image/region based registration algorithms such as Turboreg or moco (Dubbs et al., 2016), and Fourier based methods such as DFT registration or phase correlation (Pnevmatikakis et al.) for motion compensation (Thevenaz et al., 1998) Turboreg requires an initial averaged image of high enough quality in order to perform an accurate registration, while DFT in addition requires upsampling to achieve subpixel accuracy (Guizar-Sicairos et al., 2008). Moreover, more complex approaches are emerging including new algorithms for non-rigid motion compensation (Pnevmatikakis and Giovannucci, 2017). The advances in motion compensation and spike inference methods should aid in improving data quality in imaging experiments (Harris et al., 2016).

The tracking-based motion compensation introduced in this paper does not rely on an averaged image for accuracy. Instead, it extracts the position of a few reliable bright cells in every frame, tracks them in time, and corrects for the displacement without using the averaged image as reference. Tracking only the brightest cells is both more reliable and faster computationally. Subpixel accuracy increases with the pixel size of a neuron, and does not require additional computations such as upsampling. Since each tracked cell is corrected to its mean position independently, the tracking method can be expanded for use in cases where distortions of the tissue become important, e.g. near blood vessels or when imaging very large areas. Having a reference signal that is time invariant allows for calibration of the signal to noise ratio and aids in robust peak finding.

Peak finding has a second purpose in the workflow we introduce: Once image sequences are motion compensated (by any method) the peak finding algorithm is applied to the averaged images to determine the position of almost all cells visible to the eye in an unbiased way. Conversely, the subpixel accuracy motion compensation - and the ability to adjust for local deformations of the image field if needed based on the independent track of each neuron - is a strong basis for advanced spike inference and analysis tools. A novel correlation based method for example, assumes motion compensation (Pnevmatikakis et al.).

Since the analysis yields in plane motion compensation that is registered across planes for three dimensional image stack sequences, our approach provides a basis for motion compensation in all three dimensions, which promises to further enhance the signal to noise and thus enable even more accurate determination of the neuronal code.

## Acknowledgement

We acknowledge initial workflow development by Erin Marshall. Work supported by BRAIN initiative grant U01NS090569 (POK, WL) and University of Maryland Seed Support.

